# A DNA Unwinding Equilibrium Serves as a Checkpoint for CRISPR-Cas12a Target Discrimination

**DOI:** 10.1101/2023.05.16.541046

**Authors:** Jaideep Singh, Kevin G. Liu, Aleique Allen, Wei Jiang, Peter Z. Qin

## Abstract

CRISPR-associated proteins such as Cas9 and Cas12a are programable RNA-guided nucleases that have emerged as powerful tools for genome manipulation and molecular diagnostics. However, these enzymes are prone to cleaving off-target sequences that contain mismatches between the RNA guide and DNA protospacer. In comparison to Cas9, Cas12a has demonstrated distinct sensitivity to protospacer-adjacent-motif (PAM) distal mismatches, and the molecular basis of Cas12a’s enhanced target discrimination is of great interest. In this study, we investigated the mechanism of Cas12a target recognition using a combination of site-directed spin labeling, fluorescent spectroscopy, and enzyme kinetics. With a fully matched RNA guide, the data revealed an inherent equilibrium between a DNA unwound state and a DNA-paired duplex-like state. Experiments with off-target RNA guides and pre-nicked DNA substrates identified the PAM-distal DNA unwinding equilibrium as a mismatch sensing checkpoint prior to the first step of DNA cleavage. The data sheds light on the distinct targeting mechanism of Cas12a and may better inform CRISPR based biotechnology developments.

## INTRODUCTION

Clustered-Regularly-Interspaced-Short-Palindromic-Repeats (CRISPR) and CRISPR-associated (Cas) proteins provide adaptive immunity for bacteria and archaea to combat invading viruses and other mobile genetic elements (1–4). CRISPR-Cas systems have been adapted and repurposed as biotechnology tools with applications in genome engineering, transcription regulation, molecular diagnostics, cellular therapeutics, and agriculture (5–8). Class II CRISPR systems that utilize a single effector nuclease, including Cas9 and Cas12a, have found significant utilization in research and commercial settings (9). While Cas9 is the most widely utilized enzyme for genome editing, Cas12a has been adopted for genome manipulation as well as molecular diagnostic applications (10, 11).

Similar to Cas9, Cas12a uses an effector complex activated and programmed by CRISPR-encoded small RNA(s) (crRNA) to cleave double-stranded DNA (12, 13). A specific target is identified by recognition of a short protospacer-adjacent-motif (PAM) within the DNA, followed by unwinding of a segment of the DNA designated as the protospacer to form the R-loop, in which the DNA target-strand (TS) is hybridized with the single-stranded guide of the crRNA, while the DNA non-target-strand (NTS) is unwound (Fig. 1). The Watson-Crick base pairing between the RNA-guide and TS establishes a simple yet effective rule for targeting a desired genomic site and serves as the foundation for the Cas12a/Cas9-based technological evolution (14). However, it has also subjected these enzymes to undesired off-target effects, in which erroneous binding and/or cleavage occurs on off-target genomic sequences where imperfect complementarity between the RNA guide and protospacer exists (15–17). A mechanistic understanding of target discrimination is the key to combat the off-target effect, and remains an area under active investigation.

**Figure 1.**
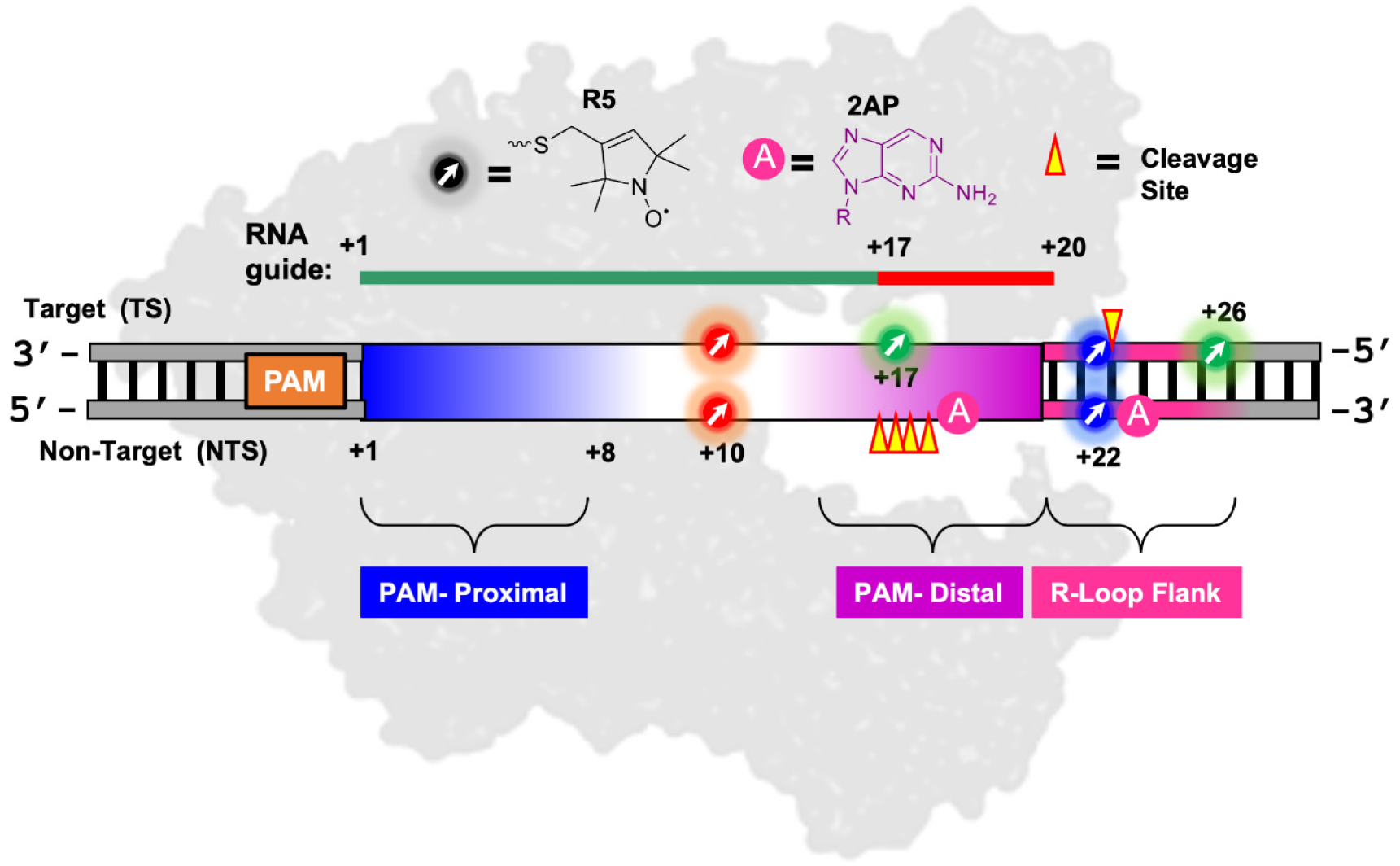
Monitoring FnCas12a induced site-specific DNA unwinding via spectroscopic labels. The FnCas12a effector is shown in grey, with the colored line representing the on-target and off-target RNA guides that vary at positions +17 to +20. The DNA substrate duplex is shown with the PAM marked by an orange box; the protospacer (PAM+1 to +20) indicated by a box with coloring varied from the PAM-Proximal (blue) to the PAM-distal segment (purple); the R-loop Flank segment beyond the RNA/DNA hybrid marked by pink lines; and cleavage sites indicated by wedges. Chemical structures of the R5 nitroxide label (indicated by the colored arrows) and the 2-aminopurine label (indicated by pink circles) are shown on top, with the corresponding labeling positions marked on the DNA. Details on DNA and crRNA guide sequences can be found in Table S1 and Table S2 respectfully.

Studies have revealed that the Cas12a-crRNA effector complex interrogates a DNA target via a series of coordinated conformational changes (18–21). Similar to Cas9, upon recognition of the PAM, Cas12a locally distorts and unwinds the base-pairs immediately adjacent to the PAM, establishing an R-Loop seed (22). Sequential unwinding of the DNA leads to formation of the full R-Loop, triggering allosteric activation and enabling DNA strand scission (18, 20, 21, 23–25). However, Cas12a differs from Cas9 in key features of effector protein, crRNA, and target DNA (12, 26). While Cas9 cleaves a DNA target using two nuclease domains in a parallel-sequential fashion (27–29), Cas12a causes double-stranded DNA breaks using a single RuvC nuclease domain following a sequential mechanism, with the NTS being cleaved first at a site located in the PAM-distal segment of the protospacer, around 14-18 nucleotides from the PAM (19). Following nicking of the NTS, the protein-RNA-DNA complex undergoes additional conformational changes, eventually enabling cleavage of the TS at a site located 1-2 nucleotides beyond the DNA/RNA hybrid within the R-loop flank (19, 30, 24, 31). Consequently, Cas12a yields a mismatch tolerance pattern that is clearly different from that of Cas9 (32), with little to no tolerance for PAM-distal mismatches (33, 34). However, while rapid progress has been made on understanding Cas12a structure (19–21) and targeting selection and discrimination (35–37), current understanding on Cas12a’s target surveillance mechanism is not as comprehensive as that of Cas9 (38, 39), and further in depth studies are needed on the structure-function relationship that governs Cas12a target selection.

Previous work has shown that in Cas9, DNA protospacer unwinding, which is directly connected with the allosteric activation of the protein, controls DNA cleavage activity and serves as a key determinant in sensing and discriminating DNA/guide mismatches (29, 38, 40, 41). Given the differences between Cas12a and Cas9 (13), in particular the mismatch sensitivity patterns (32–34), the manner of Cas12a-indcued DNA unwinding and the dynamics of the R-loop is a critical element in deciphering its mechanism of DNA target interrogation. A large number of biophysical techniques, including single molecule fluorescence and force microscopies, have been applied to explore a variety of conformational dynamics of the Cas12a ternary complex along the pathway to DNA target cleavage (20, 23, 24, 42–45), however, much remains to be learned about the R-loop conformational states of Cas12a and their roles in DNA target discrimination.

In this study, we used *Francisella novicida* Cas12a (FnCas12a) as a model system to examine Cas12a-induced DNA unwinding using a combination of site-directed spin labeling, 2-amino-purine (2AP) fluorescence, and nuclease cleavage activity measurements. The work leveraged a unique biophysical technique, site-directed spin labeling (SDSL), in which stable radicals (e.g., nitroxide spin labels) attached at specific sites of the parent molecule are utilized to obtain structural and dynamic information of the target system (46, 47). In particular, SDSL in conjunction with a pulsed Electron Paramagnetic Resonance (EPR) technique, double electron-electron resonance (DEER) spectroscopy, allows for measurements of distances and distance distributions, which provide information on dynamic equilibria between conformational states in solution. Recently, we have applied SDSL-EPR to study Cas9 (41, 48, 49), including examining site-specific Cas9-induced DNA unwinding at a resolution the of individual nucleotides (41, 49).

Results presented here reveal an inherent equilibrium between a DNA unwound state and a DNA-paired duplex-like state with an on-target Cas12a complex, in which the RNA guide fully matches the DNA protospacer. This DNA-paired/unwound equilibrium is tuned by the presence of PAM-distal RNA/DNA mismatches prior to the first step of NTS cleavage, but is no longer maintained with pre-nicked DNA substrates mimicking post-NTS cleavage states. Together, we propose a model for DNA target discrimination by Cas12a, where the DNA paired/unwound equilibrium serves as a checkpoint for licensing RuvC nuclease activity for the first step of cleavage of the NTS. The work advances our understanding on the role of DNA unwinding in directing Cas12a’s nuclease activation to achieve target discrimination and sequential strand scission.

## MATERIAL AND METHODS

### Cas12a Protein Expression and Purification

Catalytically-active FnCas12a and Catalytically-inactive dFnCas12a were expressed in *Escherichia coli* and purified following reported protocols (44, 45). Briefly, a plasmid encoding the desired Cas12a protein with a His-MBP tag at the N-terminus was transformed into Rosetta (DE3) Competent E. coli (Novagen) by heat shock. A single colony from the transformation was inoculated into Lysogeny Broth with appropriate antibiotic (50ug/mL Kanamycin) and incubated at 37°C overnight. Then, the small-scale culture was added to terrific Broth with 50 ug/mL antibiotic and incubated at 37°C until the optical density at 600 nm reached 0.5-0.6. The temperature was then reduced to 18°C, and incubation continued for 20 minutes. Overexpression was then induced by adding 200uM IPTG and shaking at 18°C for 16-20h. The cells were harvested via centrifugation and resuspended in cold lysis buffer [20mM Tris pH 8.0, 500 mM NaCl, 10 mM imidazole], with protease inhibitor (Roche). The cells were lysed at 4°C via sonication and then subjected to ultracentrifugation at 4°C to separate cellular debris. The supernatant underwent immobilized metal affinity chromatography and the pooled fractions containing His-MBP-Cas12a underwent TEV protease digestion to cleave the His-MBP tag. The crude FnCas12a was then subjected to heparin chromatography (HiTrap Heparin Hp column Cytivia). The pooled fractions were further purified by size exclusion chromatography (Cytivia S200 Increase) with 20mM Tris pH 8.0, 500mM KCl, and 1mM DTT as the running buffer, before being concentrated and stored at −80°C in storage Buffer [20mM Tris pH 8.0, 500mM KCl,1mM DTT, and 20% glycerol]. Concentrations of proteins were determined according to absorbance at 280nM with an extinction coefficient of 144 000 M^-1^ cm^-1^.

### DNA Substrate Preparation

All DNA oligonucleotides, including those with specific 2AP or phosphorothioate modifications, were synthesized via solid-phase chemical synthesis and obtained commercially (Integrated DNA Technologies, Coralville, Iowa). Complete information of DNA substrate sequences is described in Supporting Information Table S1. DNA concentrations were determined by measuring the absorbance at 260nm and using the extinction coefficient values provided by the vendor.

Formation of the target DNA duplex was carried out via mixing a 1:1 molar ratio of the desired target and non-target strand in annealing buffer (50mM Tris, pH 7.5, 100mM NaCl) and incubating overnight at room temperature. The annealed duplexes were purified by size exclusion chromatography (SEC) and buffer-exchanged into 1X CRISPR reaction buffer (20mM Tris, pH 7.5; 150mM KCl, 5% glycerol, and 5mM MgCl_2_) and stored at −20°C until measurement.

### crRNA Preparation

The crRNA’s were prepared by T7 *in vitro* transcription using a T7 top-stranded primer and a single-stranded DNA oligonucleotide template (see Table S2 for detailed sequences). Transcription reactions were carried out following reported procedures (44). The crude transcription products were purified via denaturing polyacrylamide gel electrophoresis and redissolved in ME buffer (10 mM MOPS, pH 6.5, and 1mM EDTA) and stored at −20°C. RNA concentrations were determined based on the absorbance at 260nm, with the molar extinction coefficients estimated by ɛ = number of nucleotide × 10,000 M^−1^·cm^−1^. crRNA sequences are listed in Table S3.

### Preparation of dCas12a Ternary Complexes for Spectroscopy Measurements

Ternary complexes were assembled in 1X CRISPR reaction buffer with the dCas12a/crRNA//DNA molar ratio being 1:1.2:1.1. To assemble the complex, first the crRNA was refolded via heating at 95°C for 1min in ME buffer and then cooled at room temperature for 5 min. The crRNA was adjusted to an appropriate amount of salt to match the 1X CRISPR reaction buffer and then mixed with the dCas12a protein. The dCas12a/crRNA mixture was incubated at 37°C for 10 minutes to form the effector complex, and then the appropriate amount of DNA duplex was added. The dCas12a/crRNA/DNA mixture was incubated at 37°C for an additional 10 minutes before purification using SEC in 1X reaction buffer. SEC fractions containing ternary complexes with 2AP modified DNA were pooled and used directly for 2AP fluorescence measurements. Ternary complexes containing spin-labeled DNA were further processed as described below.

### 2-amino-purine (2AP) Fluorescence Measurements

2AP measurements were conducted using a Molecular Devices SpectraMax® iD5 plate reader. 100uL of purified 2AP-modified duplex DNA or dCas12a ternary complex was transferred to sample wells in a black Corning® 384 well flat bottom polystyrene plate (Mfr No. 35766). Utilizing the onboard monochromator, the excitation was fixed at 315 nm with the fluorescence emission maxima collected at 368nm. In parallel, absorbance measurements on the corresponding samples were obtained on a DU800 UV−Vis spectrometer (Beckman Coulter, Fullerton, CA). Following a prior report (41), the background-corrected emission at 368nm (F_368_, obtained by subtracting the corresponding buffer emission) was normalized by the 260nm absorbance (A_260_) of the same sample to obtain:

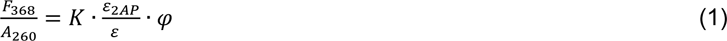

where ε is the extinction coefficient of the particular sample, ε_2AP_ is the extinction coefficient of 2AP at 315 nm, φ is the quantum yield of 2AP, and K is a constant independent of the sample concentration. Furthermore, one can normalize the 2AP quantum yield of a given sample to that of the corresponding free duplex as:

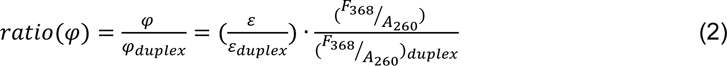

*ratio (*𝜑), which specifically reports on the 2AP quantum yield while minimizing the impact of local DNA sequence variations, was used to access DNA unwinding.

### Site-Directed Spin Labelling

Site-specific labeling with the R5 nitroxide label was conducted following a previously described protocol (50, 51). Briefly, crude phosphorothioate-modified DNA oligonucleotide was mixed with an acetonitrile/water (50%/50% by volume) solution containing a R5 precursor [3-Iodomethyl-1-oxyl-2,2,5,5-tetramethylpyrroline (Toronto Research Chemicals, cat. no. I709500)] (60-100mM) and 100 mM MES (pH 5.8). The mixture was incubated at room temperature for 12-24 hours. The labeled oligonucleotide was then purified into sterile deionized water via application of a desalting spin column (Thermo Scientific Zeba 7K MWCO, cat. No. 89883). Note that follow-up SEC purification of the DNA duplex or the ternary complex served as an additional step to remove unattached nitroxides. Labeling efficiency was assessed by a previously reported spin counting procedure (52) and was generally >90%. The R5-labled oligonucleotide was stored at −20°C until DNA substrate preparation.

### Continuous Wave EPR Measurements

Spin-labeled samples (10-50uM) were transferred to round glass capillary tubes (0.6 mm ID, 0.8 mm OD; Vitrocom, Inc.) for measurement. The continuous-wave EPR (cw-EPR) spectra were measured utilizing a Bruker EMX spectrometer housing an ER4119HS cavity. For spectral acquisition, the incident microwave power was 2mW, and the field modulation was 2G at a frequency of 100 kHz. Each spectrum was acquired with 512 points, corresponding to a spectral range of 100 G, then corrected for background baseline, and intensity normalized following reported protocols (52).

### Double Electron-Electron Resonance (DEER) Spectroscopy

Doubly R5-labeled samples were buffer exchanged to 1X CRISPR reaction buffer dissolved in deuterium oxide via a 100 kDa spin filter concentrator, then combined with the appropriate volume of deuterated glycerol to produce DEER samples in 30% (v/v) glycerol. Each DEER sample was approximately 30uL, with the sample concentration estimated to be 60-80 uM. The samples were loaded into quartz EPR capillary tubes and flash frozen with liquid nitrogen immediately before measurement.

DEER measurements were carried out at 50 K on a Bruker ELEXSYS E580 X-band spectrometer with an ER4118-MS3-EN resonator. A dead-time free four-pulse scheme was used (53), with the pump pulse frequency set at the center of the nitroxide spectrum and the observer frequency being approximately 70 MHz lower. The observer p pulse was 32 ns. The pump π pulse, optimized using a nutation experiment, was set between 8 and 16 ns. The delay between the observer π/2 and π pulses was set to 420 ns to minimize deuterium ESEEM. The video bandwidth was fixed at 20 MHz. The shot repetition time was set between 1,000 and 1,400 μs. Accumulation time in each measurement was approximately 20–24 h, with 100 shots per point.

Distance distributions from the DEER data were obtained using Consensus Deer analysis, an analysis package accompanied in the DEERAnalysis software kit (54, 55). Modeling of inter-R5 distance distributions was performed using ALLNOX (56, 57). The input structure for the free DNA was a linear B-form duplex built with the P3 sequence using Avogadro (58). The input structure for the ternary complex was a FnCas12a crystal structure (PDB 6i1k) showing unwound DNA (19). Modeling used previously reported parameters (57), except that the torsion angles were varied with a step size of 60°. Modeling yielded the ensemble of pairwise distances between the nitrogen atoms of the pyrroline of the allowed R5 conformers (N-N), from which the average distance was computed (56, 57).

### DNA Cleavage Assay

DNA substrate duplexes containing terminal fluorescent labels were purified by SEC into 1 X CRISPR buffer. For a given reaction, a Cas12a/crRNA effector complex (200nM) was assembled with a protein/crRNA molar ratio of 1:1.25 and incubated in 1X CRISPR reaction buffer for 10 min at 37°C. The desired DNA substrate (25nM) was then added. The reaction was allowed to proceed at 20°C for the desired time, before being stopped via addition of denaturing gel loading buffer (95% formamide, 20mM EDTA, 0.025% SDS) with the volume ratio between the dye and the sample being 1:4. The cleavage products and the precursor were resolved via a denaturing polyacrylamide gel. The resulting gels were imaged using a ChemiDoc MP system (Bio-Rad) using the appropriate fluorescein imaging settings. The band intensities of the DNA species and background were quantified using ImageJ (1.47). The fraction of precursor for each timepoint was calculated as:

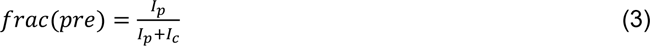

with I_p_ being the background corrected uncleaved precursor band intensity, and I_c_ being the background corrected intensity of the cleaved product. Cleavage experiments were performed in triplicate, and the average *frac(pre)* was plotted for each timepoint with the standard deviation shown as error bars. When appropriate, the time dependence of average *frac(pre)* was fit to a single-exponential decay using an in-house program written in Matlab:

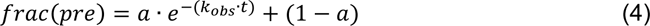

with k_obs_ representing the rate constant, and *a* representing the active fraction of the precursor.

## RESULTS

### A DNA Duplex-Like Paired State Coexists with the Unwound State in the On-Target Ternary Complex Prior to DNA Cleavage

To evaluate Cas12a mediated DNA unwinding using SDSL, we constructed a model DNA substrate duplex termed P3, and attached R5 nitroxide labels onto desired specific backbone positions (Fig. 1; Table S1). The labeled duplex was assembled into a ternary complex with a catalytically-inactive FnCas12a/crRNA effector (dCas12a). The dCas12a ternary complex was purified (Fig. S1A) then subjected to DEER measurements. Control measurements showed only minor impact on DNA cleavage rates due to spin-labeling, indicating the R5 labels did not significantly perturb the native folding and function of the Cas12a ternary complex (Fig. S2).

DEER measurements were first carried out with a pair of R5 labels attached at backbone positions directly across the duplex at the PAM+10 protospacer positions (Fig. 2A and B), which are located in the middle of the protospacer (Fig. 1). With the free (unbound) DNA duplex, the observed dipolar evolution trace measurement showed oscillations with fast decay and yielded a distance distribution showing a single population centered at 26Å (Fig. 2A). This is consistent with previously reported studies of R5-labled DNA duplexes (50). With an on-target ternary complex, where the DNA was bound to a dCas12a effector containing an RNA guide that fully matches the DNA protospacer, the measured dipolar evolution showed oscillations containing features of both fast and slow decay (Fig. 2B). The resulting distance distribution showed two well-defined major populations, one centered at 28Å and another centered at 41Å (Fig. 2B). Modeling using a reported dCas12a on-target complex crystal structure (PDB 6i1k), which shows unwinding of the DNA protospacer (19) resulted in a predicted average inter-R5 distance of 43.5Å at PAM+10 (Fig. 2B, Model). Considering the 2- 3Å range of variation in average inter-R5 distances in the modelling (56, 57), the predicted PAM+10 distance on the 6i1k structure showing DNA unwinding matches very well with the measured long-distance population (Fig. 2B), leading to the conclusion that the observed long distance represents the DNA unwound state.

**Figure 2.**
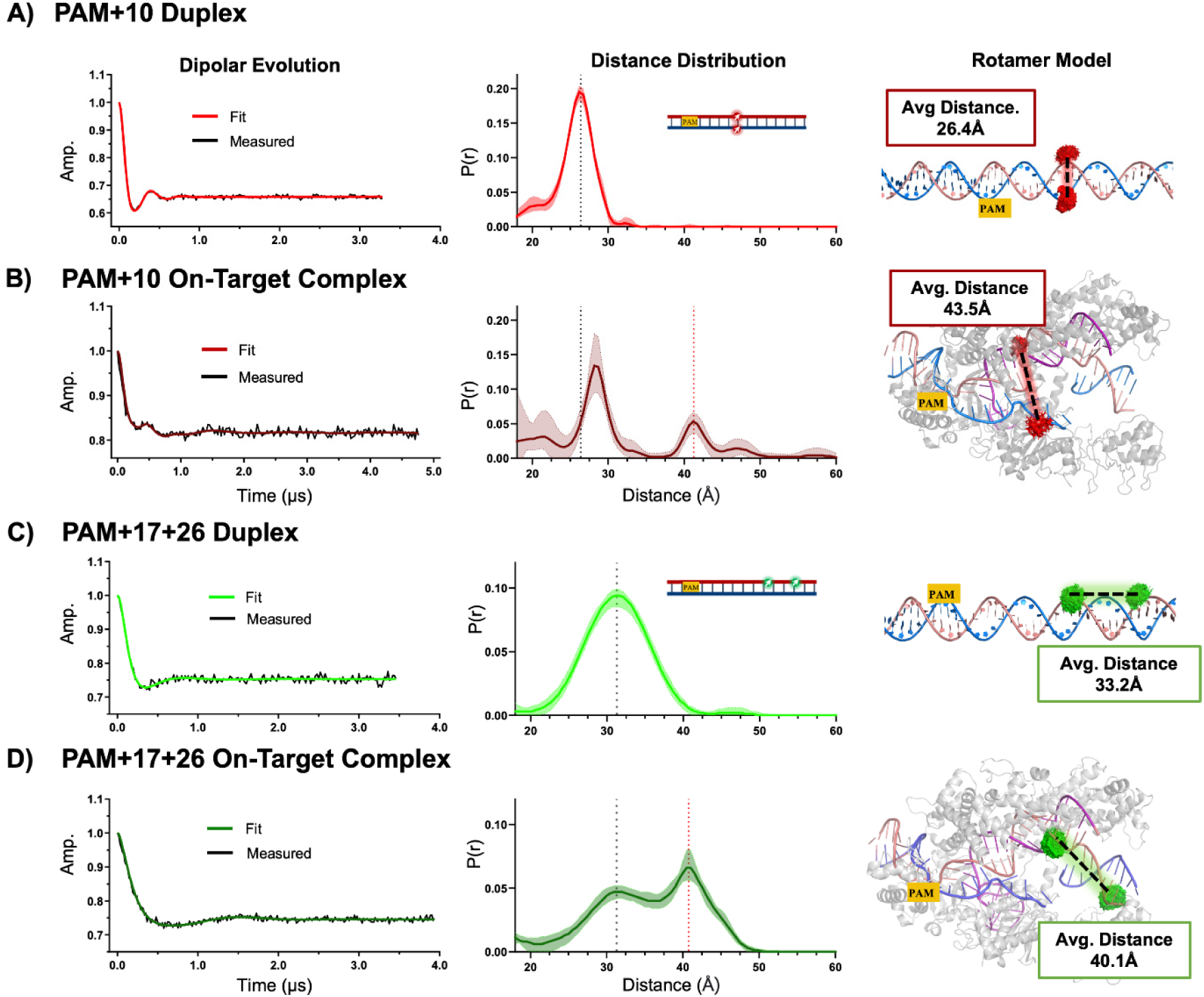
A DNA duplex-like paired state coexists with the unwound state in the on-target ternary complex prior to DNA cleavage. Distance measurements are shown for the PAM+10 (panels (A) and (B)) and the PAM+17+26 (panels (C) and (D)) datasets obtained with intact NTS strands. For each data set, shown on the left is the DEER dipolar evolution trace; middle is the resulting distance distribution with the error contour represented by the shaded band; and right is the rotamer modeling obtained with ALLNOX. To aid comparison, the distance distributions include a black dashed line to mark the maximum distance measured from the corresponding free DNA duplex, and a red dashed line to mark the maxima of the long-distance populations measurand on the complex samples, which represent the DNA in an unwound state. Additional details can be found in Figures S3, S4, and Table S4.

It is interesting that the DEER data reveal a significant 28Å-population at PAM+10 within the on-target complex (Fig. 2B). This distance is within 2Å of the inter-R5 distance value predicted using a uniform B-form duplex model (Fig. 2A, Model) as well as that measured with the corresponding free DNA (Fig. 2A). It indicates that the corresponding population undergoes very little, if any, strand separation (i.e., unwinding), and therefore is assigned as a “duplex-like” state with the DNA remaining paired at the PAM+10 position. Note that control measurements confirmed that the large majority of the R5-labeled DNA was bound within the dCas12a complex (Fig. S1B). Given the significant presence of the 28Å population (Fig. 2B), it is impossible that it arose from DNA dissociated from the ternary complex. In addition, R5 labels attached at the PAM+10 sites, which are located at the middle of the protospacer, likely incur a fair number of contacts to dCas12a and crRNA within the ternary complex, as indicated by cw-EPR measurements (Fig. S5). This may contribute to the difference between the observed 28Å distance within the complex vs. that measured with the free DNA (26Å, Fig. 2A). As such, the PAM+10 DEER data (Fig. 2B) reveal that the bound DNA exists in equilibrium between a duplex-like state and an unwound state.

Although control measurements showed only minor impacts on DNA cleavage rates due to R5 labeling (Fig. S2), one cannot completely exclude the possibility that R5 labels presented at PAM+10 sites bias DNA unwinding to some degree. To further examine the possible simultaneous presence of the duplex-like and unwound states in the on-target ternary complex, we prepared a P3 construct with R5 attached at the TS PAM+17 and TS PAM+26 positions (Fig. 1, Table S1), and measured this P3+17+26 distance to examine the target-strand conformation (Fig. 2C and 2D). DEER measurements with the free duplex revealed a population with a distance distribution centered at 31Å (Fig. 2C), which agrees quite well with the model-predicted distance (33.2Å, Fig. 2C, model). When bound to the on-target dCas12a ternary complex, the DEER measured distance profile showed a clear bimodal distribution (Fig. 2D). A major population was centered at 41Å. This matches the predicted value of 40.1 Å obtained using the 6i1k crystal structure (Fig. 2D, Model), which shows DNA unwinding at the PAM+17 position and kinking of the R-loop flank segment beyond the 20-base-pair RNA/DNA hybrid. This population therefore represents the unwound DNA. Importantly, the other major population is centered at 31Å, which matches exactly with the distance measured in the free linear duplex (Fig. 2C), and therefore was assigned as the duplex-like state representing the paired DNA. As such, the data reveal that the PAM-distal segment of the DNA target strand exists in equilibrium between an unwound state and a duplex-like paired state. We note that the PAM+10 data set seems to show a higher percentage of the duplex-like population than that observed in the PAM+17+26 data set (Fig. 2B and 2D), which may reflect site-specific variations of R5’s impact on the bound DNA conformations. None the less, the clear presence of the bimodal distribution observed with the PAM+17+26 data set unambiguously support the conclusion that within the on-target complex, an equilibrium of the unwound and duplex-like paired states exists at the protospacer segment distal from the PAM.

### The PAM-Distal DNA Paired/Unwound Equilibrium Serves as a Checkpoint to License the First-Step NTS Cleavage

Next, we examined how DNA unwinding is impacted by mismatches between the DNA target and the RNA guide. We focused on mismatches at the PAM-distal positions, as there is a clear difference in mismatch sensitivity at this segment between Cas12a and Cas9 (32, 33). A P3 off-target crRNA (Table S3) was synthesized to assemble an off-target ternary complex with dCas12a and the P3 DNA, with the PAM+17 to +20 positions (i.e. the most PAM- distal portion of the RNA/DNA hybrid) all containing RNA/DNA mismatches. DEER measurements were carried out for this P3 off-target ternary complex with the R5-labeled P3+17+26 DNA. The resulting distance distribution showed only one population with the maxima located at 31Å (Fig. 3A) that is nearly identical to that measured from the free duplex (Fig. 3B). Furthermore, comparing to the P3 on-target complex, the off-target complex clearly lost the 41Å population that represents the unwound DNA (Fig. 3B). The DEER data therefore revealed that the four mismatches disrupt the DNA unwinding equilibrium -- the DNA unwound state is lost, and the DNA remains in the duplex-like paired state in the ternary complex.

**Figure 3.**
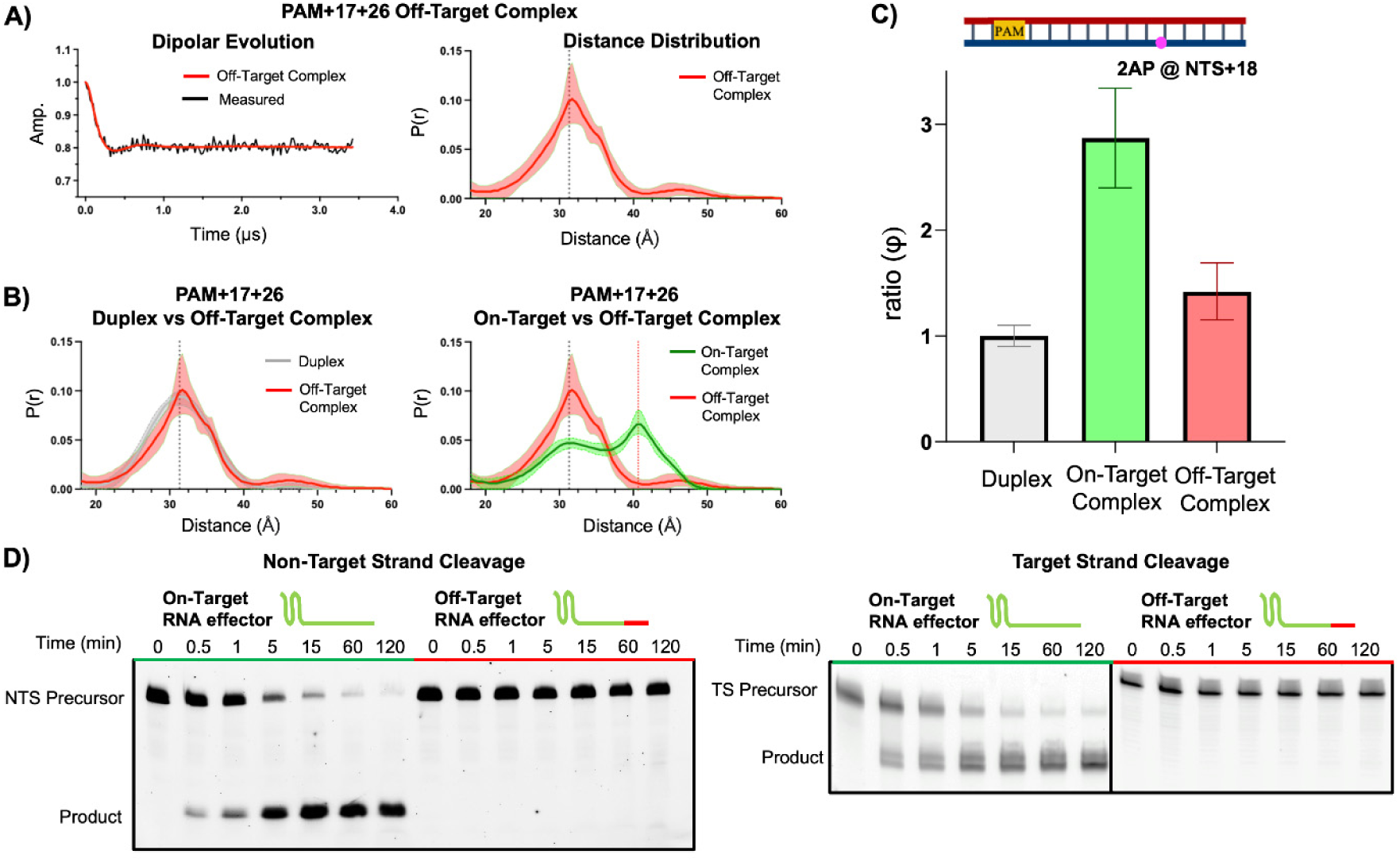
The PAM-distal DNA unwinding equilibrium serves as a checkpoint for the first step of NTS cleavage. (A) Distance measurements on the PAM+17+26 intact duplex bound to an off-target effector that contains 4 consecutive protospacer mismatches at positions PAM+17-20. (B) Comparison of the PAM+17+26 distance distributions between the free duplex and the off-target complex (left) and the on-target and off-target complexes (right). To aid comparison, the distance distributions include a black dashed line to mark the maxima measured from the corresponding DNA duplex, and a red dashed line to mark the maxima of the long-distance (unwound DNA) population measurand on the complex samples. Additional details can be found in Figures S3, S4, and Table S4. (C) Normalized fluorescence quantum yield (ratio(ϕ)) obtained with the 2AP incorporated at the NTS PAM+18 position of the P3.2 construct. Additional details can be found in Table S5. (D) Representative gels showing time course measurements of NTS (left) and TS (right) cleavage by FnCas12a.

Further investigations on PAM-distal DNA unwinding were carried out using a 2AP fluorescence assay developed in our prior Cas9 study (41). A slightly modified DNA substrate, P3.2-2AP (Table S1), was generated to substitute a 2AP at the NTS PAM+18 position. P3.2- 2AP was assembled into either an on-target complex (i.e., RNA/DNA match from PAM+1 to PAM+20) or an off-target complex (i.e., mismatches at PAM+17 to PAM+20). Following our prior work (41), *ratio (*𝜑) (eq.2), which reports on the 2AP quantum yield and serves as a sensor of the state of stacking of the DNA base, was obtained to access DNA unwinding. For the on-target complex, the measured *ratio (*𝜑) value was 2.87±0.47, which is clearly higher than that obtained for the free duplex (1.00±0.10) (Fig. 3C, Table S5). This is consistent with the expected DNA unwinding at PAM+18, as unwinding would reduce DNA base stacking, thus increasing 2AP quantum yield. For the off-target complex, the *𝑟𝑎𝑡𝑖𝑜(𝜑)* value was 1.47±0.27, which increases only slightly compared to that of the duplex (Fig. 3C, Table S5). This indicates little DNA unwinding. As such, the 2AP measurements strongly support the conclusions drawn from the DEER data, that the PAM-distal mismatches push the system back to the duplex-like paired state, while the DNA unwound state is not populated.

We further carried out in vitro DNA cleavage measurements using catalytically-active FnCas12a to examine the functional consequences of perturbing the equilibrium in Cas12a induced DNA unwinding. Under saturating enzyme concentrations, a complex containing the on-target RNA cleaved the NTS to completion within 120 minutes, while a complex containing the off-target RNA resulted in no observable products during the same time period (Fig. 3D), indicating a loss of the ability to cleave NTS. Given that with Cas12a, NTS cleavage is required for TS cleavage, the off-target complex showed no cleavage product for the TS, while the on-target complex cleaved the TS to completion as expected (Fig. 3D). Furthermore, control experiments confirmed that the DNA substrate was completely bound by either the on-target or the off-target complex (Fig S6).

Taken together, with four RNA/DNA mismatches at the PAM-distal segment, pairings between the DNA TS and NTS dominate, and the system is trapped in the duplex-like DNA paired state. This results in a complete loss of the DNA unwound state, preventing the first-step cleavage of the NTS and subsequently the cleavage of the TS. As such, the DNA paired/unwound equilibrium acts as checkpoint, with occupancy of the unwound state serves as a license to allow progression to DNA strand scission.

### Nicking the Non-Target-Strand Alters DNA Conformation at the Target-Strand Cleavage Site

Prior studies have shown that following DNA unwinding and R-loop formation, FnCas12a first nicks the NTS at a site located close to PAM+18 and has trimming activity on the NTS up to PAM+15 (19, 30) (Fig. 1). The activated RuvC nuclease then continues to cleave TS at/near PAM+22, which is located beyond the 20-base-pair RNA/DNA hybrid (Fig. 1), to generate a double-stranded DNA break (19, 30). It is believed that conformational changes of DNA must occur at the target-strand cleavage site after NTS nicking but prior to TS cleavage, however, details remain largely unknown (24, 30, 45). In the SDSL and 2AP measurements described above, dCas12a and P3 duplexes containing an intact NTS and TS (Fig. 2 and 3) were used to investigate DNA unwinding prior to the first catalytic step, i.e., NTS nicking by Cas12a. To further investigate DNA conformational changes at the target-strand cleavage site in solution beyond the first catalytic step, we developed P3-nicked DNA constructs, in which the intact TS oligonucleotide (i.e, P3a, Table S1) is hybridized with two oligonucleotides to generate a fully paired duplex with a nick at/near the NTS cleavage site (Table S1). The nicked substrates were reconstituted with dCas12a and various crRNAs to mimic post NTS-nicking ternary complexes that are progressing towards TS cleavage.

SDSL and 2AP measurements were first conducted to investigate DNA conformation at the TS cleavage site. A pair of R5 nitroxides were attached at cross-strand positions at PAM+22, and the labeled DNAs were bound with the fully-matched crRNA effectors to assemble on-target ternary complexes (Fig. 4). With the P3 construct containing an intact NTS, the on-target complex gave a dipolar evolution trace showing oscillations with a fast decay, and the resulting distance distribution showed a dominant population centered at 26Å (Fig. 4A, complex). A minor peak was also observed with the center being at 20Å (Fig. 4A, complex), although given its large error contour it is not clear whether this represents an artifact or a true minor population. The major 26Å population agrees well with the predicted value of 28.2 Å obtained using the 6i1k structure (Fig. 4A, complex model), which is representative of an intact NTS in an on-target complex. Furthermore, PAM+22 measurement on the free intact DNA yielded a distance distribution with one population centered at 26Å (Fig. 4A, duplex), and cw-EPR measurements indicated that the DNA local environment at NTS+22 is similar to that of a free duplex without incurring contacts to the Cas12a effector (Fig. S5C). Together, SDSL indicated that prior to NTS nicking, in solution the target-strand cleavage site (i.e., PAM+22) remains at the duplex-like DNA paired state. In addition, with 2AP incorporated at NTS PAM+23, which is directly adjacent to the target-strand cleavage site, the complex gave a *ratio (*𝜑) value of 0.95±0.04 (Fig. 4C), which is comparable to the free duplex (Fig. 4C, intact) and indicates the base remains well stacked. Overall, prior to NTS nicking, while the protospacer segment exists in equilibrium between the unwound and duplex-like paired states, the target-strand cleavage site, which is at the R-Loop flank downstream of the protospacer, remains in a duplex-like state.

**Figure 4.**
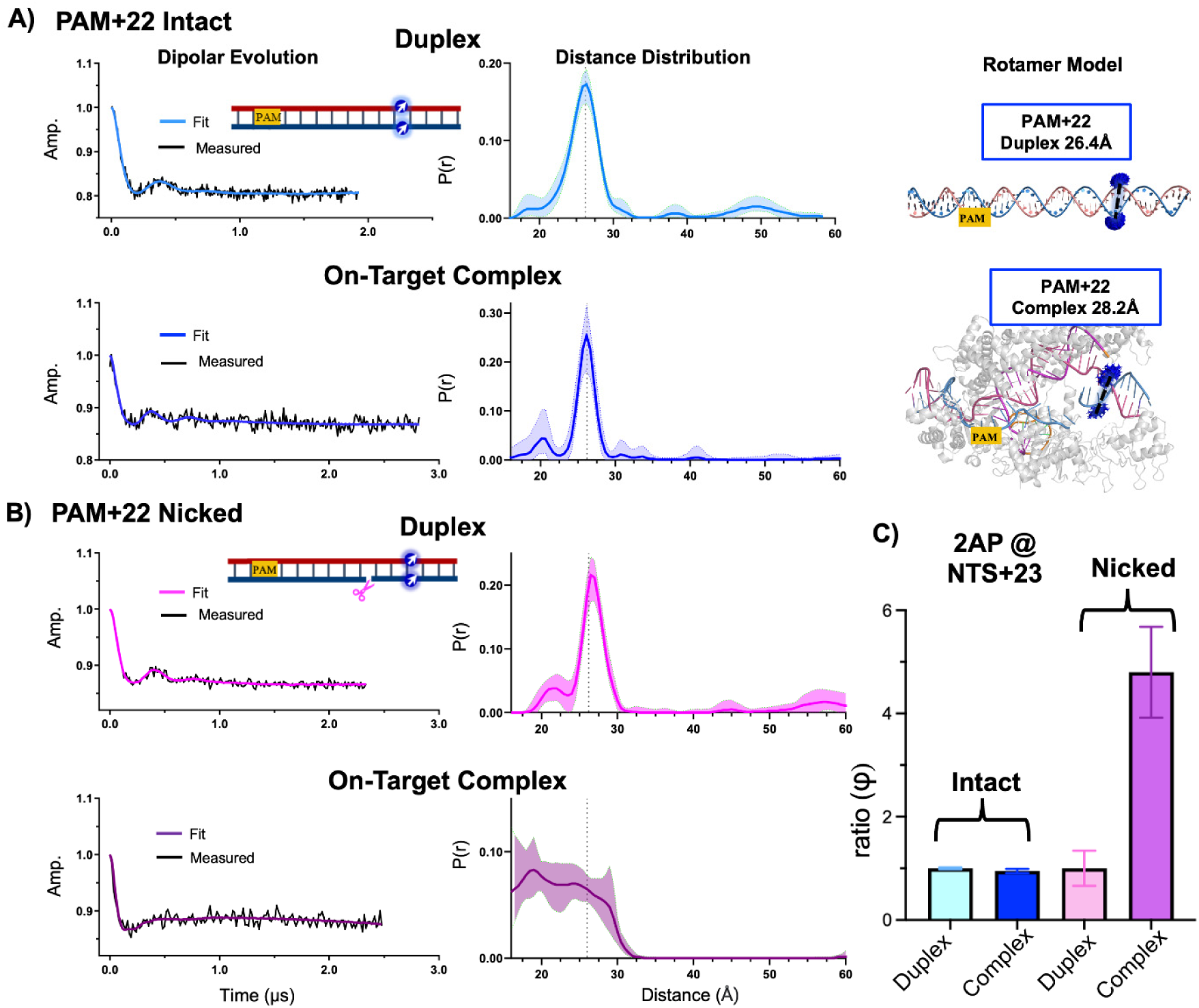
Non-target strand nicking alters DNA conformation at the target-strand cleavage site. (A) Distance measurements of intact DNA labeled across the duplex at the target strand cleavage site (PAM+22). Shown on the left is the DEER dipolar evolution trace; middle is the resulting distance distribution with the error contour represented by the shaded band and the maximum distance measured from the free intact DNA duplex marked by a dotted line; and right is the rotamer modeling obtained with ALLNOX. (B) Distance measurements of the pre-nicked P3-n15 duplexes labeled at PAM+22, with dipolar evolution traces shown on the left and the resulting distance distribution shown on the right. (C) Fluorescence quantum yield measurements of 2AP incorporated at NTS PAM+23 for the intact P3 and pre-nicked P3-n15 constructs. Additional details on DEER measurements can be found in Figures S3, S4, and additional detail on 2AP measurements can be found at Table S5.

When the P3 construct with a NTS nick at PAM+15 (P3-n15, Table S1) was assembled into the on-target complex, the PAM+22 DEER data differed clearly from that of the intact P3 (Fig. 4). The free P3-n15 construct behaved as a typical duplex: the dipolar evolution trace showed oscillations, and the resulting distance distribution showed one major population centered at 26Å (Fig. 4B, duplex). DEER signal, which require the simultaneous presence of R5 at the TS and the nicked NTS, was also observed from bound P3-n15 (Fig. 4B, complex), indicating the nicked strand remained associated with the ternary complex. However, oscillations were not observed in the dipolar evolution trace of the bound P3-n15 (Fig. 4B, complex), and the resulting distance distribution showed a broad profile with no definable populations (Fig. 4B, complex). This highly heterogeneous PAM+22 distance profile likely reflects a number of aspects of conformational changes that occur upon NTS nicking. The DNA segment may move into multiple new positions with respect to the protein and/or RNA, resulting in heterogeneous contacts between the R5 labels and the parent complex that alter inter-R5 distances. This is supported by the observed changes in the cw-EPR spectra between the nicked and intact PAM+22 NTS complex (Fig. S5C). More importantly, with these movements, the DNA segment likely experiences distortion/unwinding. This is supported by 2AP fluorescence data. With a 2AP substituted at NTS PAM+23, the P3-n15 construct gave a *ratio (*𝜑) value of 4.80±0.88 (Fig. 4C, nicked). This is substantially higher than that of the free nicked duplex (Fig. 4C, nicked), and indicates significant unstacking of the base at the TS cleavage site. Overall, with the NTS nicked, the SDSL and 2AP data reveal alterations of DNA conformation at the TS cleavage site in solution. This likely reflects the series of complex transformations that occur as the system transitions toward target-strand cleavage, although discerning details about these transitions are beyond the capability of the spectroscopic tools used in this work.

### The PAM-Distal DNA Paired/Unwound Equilibrium Checkpoint Functions Prior to Nicking of the Non-Target Strand

To further examine PAM-distal DNA unwinding and its role in target discrimination after the first-step of NTS nicking, we measured the PAM+17+26 distances using a P3 construct with a nick at NTS 18 (P3-n18, Table S1). With the free duplex, the distance distribution obtained for the nicked duplex is centered at the same value as that of the intact duplex, and it shows a slightly larger width that is consistent with an increasing flexibility due to nicking (Fig. 5A, duplex). However, with the on-target complex, the dipolar oscillation trace with P3-n18 showed decay with no oscillation, and the resulting distance distribution was very broad (Fig. 5A, on-target complex). Different from that of the intact construct (Fig. 5A and B), the nicked on-target complex distance distribution showed no discernible discrete populations (Fig. 5A, on-target complex). The profile showed highest occupancy at 35 Å, which is slightly longer that the 31Å maxima of the free nicked duplex (Fig. 5A). A fair amount of occupancy was also represented at the 41Å segment that was identified as the unwound/bent conformation (Fig. 5A on-target complex). The data indicate that alteration of DNA conformation upon nicking the NTS is not only confined to the TS cleavage site (i.e., PAM+22 & PAM+23, Fig. 4), but also to the R-Loop flank downstream of the protospacer. The system no longer occupies, in an equilibrium fashion, the discrete unwound state and the duplex-like paired state (Fig. 5B, on-target complex), but instead samples a larger number of conformations. Considering that the complex was constituted with dCas12a which is unable to execute DNA strand scission, it is likely that the system samples continuously through multiple states along the reaction pathway, resulting in the observed highly heterogeneous distance distribution profile (Fig. 5A, on-target complex).

**Figure 5.**
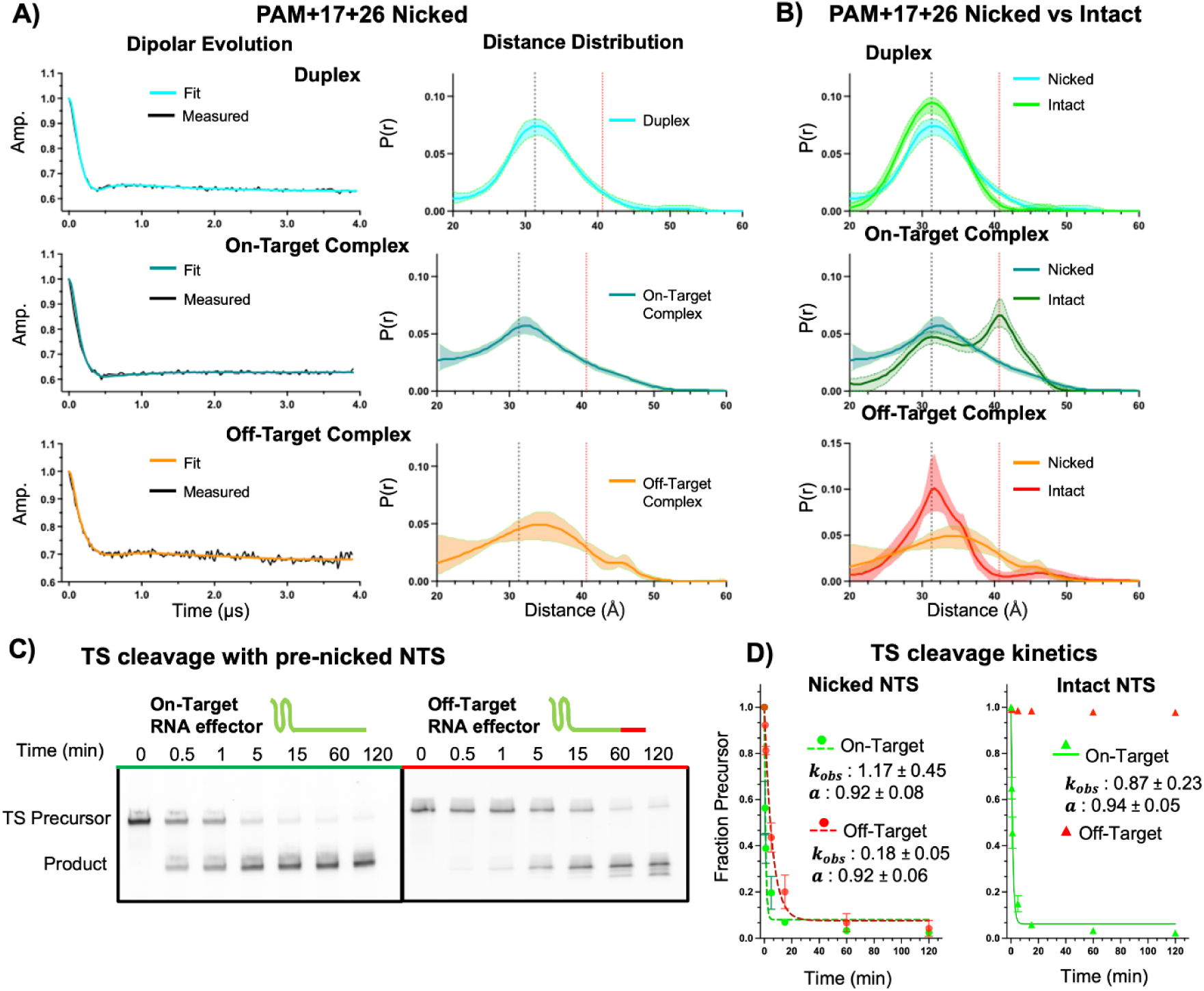
The PAM-distal DNA unwinding equilibrium is lost upon NTS nicking. (A) Distance measurements obtained with the pre-nicked P3-n18 constructs labeled at PAM+17+26. Shown on the left is the DEER dipolar evolution trace; and on the right is the resulting distance distribution with the error contour represented by the shaded band. For comparison, on the distance distributions a black dotted line marks the maxima of the free nicked DNA duplex, and a red dotted line marks the maxima of the unwound DNA measured from the intact PAM+17+26 sample (see Figure 2D). Additional details can be found in Figures S3 and Table S4. (B) Comparisons of distance distributions obtained for the NTS pre-nicked and intact constructs. (C) Representative gels showing TS cleavage by Cas12a on the pre- nicked P3-n18 substrates; (D) Fitting analysis to obtain TS cleavage rate constants. Data shown in Figure 5C are used to obtain values for the NTS pre-nicked substrate. Data shown in Figure 3D are used to obtain values for the NTS intact substrate with the on-target complex, while values for the off-target complex is not fit due to the lack of reaction progression. Note that FnCas12a exhibits imprecise NTS cleavage and trimming activity between positions NTS+15 to NTS+18 (19, 30). While P3-n18 and P3-n15 (Figure 4) place the nick at different NTS sites, they both mimic the post-NTS-cleavage DNA, and serve as the proper constructs to study progression towards TS cleavage by Cas12a.

When the P3-n18 construct was assembled into the off-target complex, the measured PAM+17+26 distance distribution profile was again very broad and without discernible discrete populations (Fig. 5A, off-target complex). The maximum of the off-target complex differs from that of the on-target complex (Fig. 5A), indicating that the RNA/DNA mismatches do impact the DNA conformation at the R-loop flank segment. Furthermore, compared to the intact off-target complex (Fig. 5B, off-target complex), the nicked off-target complex showed clearly higher relative occupancy at the long-distance region that includes the 41Å segment that is identified as the unwound/bent conformation. This indicates that with NTS pre-nicked, the off-target complex is able to sample through the conformational states required for progressing toward TS cleavage. Indeed, an active Cas12a with the off-target RNA cleaved the TS of the nicked P3-n18 substrate to completion (Fig. 5C), which is drastically different from that of the intact substrate (Fig. 5D). Furthermore, off-target cleavage with the nicked P3-n18 was 6.5 times slower than that of on-target cleavage (Fig. 5D). This indicates conformation at the R-loop flank segment remains different between the on-target and off-target complexes, and is consistent with the difference in the observed PAM+17+26 distance distribution profiles (Fig. 5A).

With the P3-n18 construct, both on-target and off-target complexes gave broad PAM+17+26 distance distribution profiles showing no discrete populations (Fig. 5A), indicating that the system is sampling a number of conformations instead of occupying the discrete DNA paired and DNA unwound states. In parallel, the nicked construct gave a much smaller off-target rate reduction in TS cleavage (Fig. 5D). Together, this indicates that the equilibrium between the duplex-like paired state and the DNA unwound state serves as a checkpoint to discriminate against RNA/DNA mismatches prior to NTS nicking, but is no longer effective after the first-step DNA strand scission.

## DISCUSSION

Based on the data presented above, we propose a model in which an equilibrium between a duplex-like paired state and a DNA unwound state serves as a check-point to interrogate PAM- distal RNA/DNA mismatches, which licenses Cas12a for the first-step of NTS cleavage (Fig. 6). As the Cas12a effector engages an intact DNA duplex, PAM-recognition followed by R-loop formation at the PAM adjacent 5-8 base-pair segment achieves binding of the DNA to the ternary complex. For an on-target DNA, propagation of DNA unwinding and R-loop formation from the mid-protospacer to the PAM-distal segment establishes the fully unwound conformation, which has been previously observed in crystal structures (19) and is captured by DEER measurements in solution (Fig. 2). Interestingly, the DEER data revealed another conformation, with the bound DNA showing distance distributions matching that of a paired duplex (Fig. 2B and C). We note such a duplex-like paired DNA conformation at PAM-mid to distal segment is also observed in newly reported cryo-EM structures (21). Importantly, the DEER data clearly show that both states are present simultaneously (Fig. 2 B & C), indicating that they co-exist in solution at equilibrium (Fig. 2 and Fig. 6). This conclusion is consistent with several prior studies showing that the Cas12a R-loop is reversible and can sample multiple states in a dynamic equilibrium(24, 35, 42–45). We also note that DEER and 2AP data obtained on FnCas12a here show that with intact NTS, the TS cleavage site (i.e., PAM+22) remains at the duplex-like paired state with little DNA distortion (Fig 4A & C). While this is consistent with the reported FnCas12a crystal structure (19) (Fig. 4A), it contrasts with other reports with LbCas12a (24, 30). This may reflect variations in R-loop dynamics and mismatch discriminations among different Cas12a orthologs reported in a number of prior work (30, 33, 37, 43).

**Figure 6.**
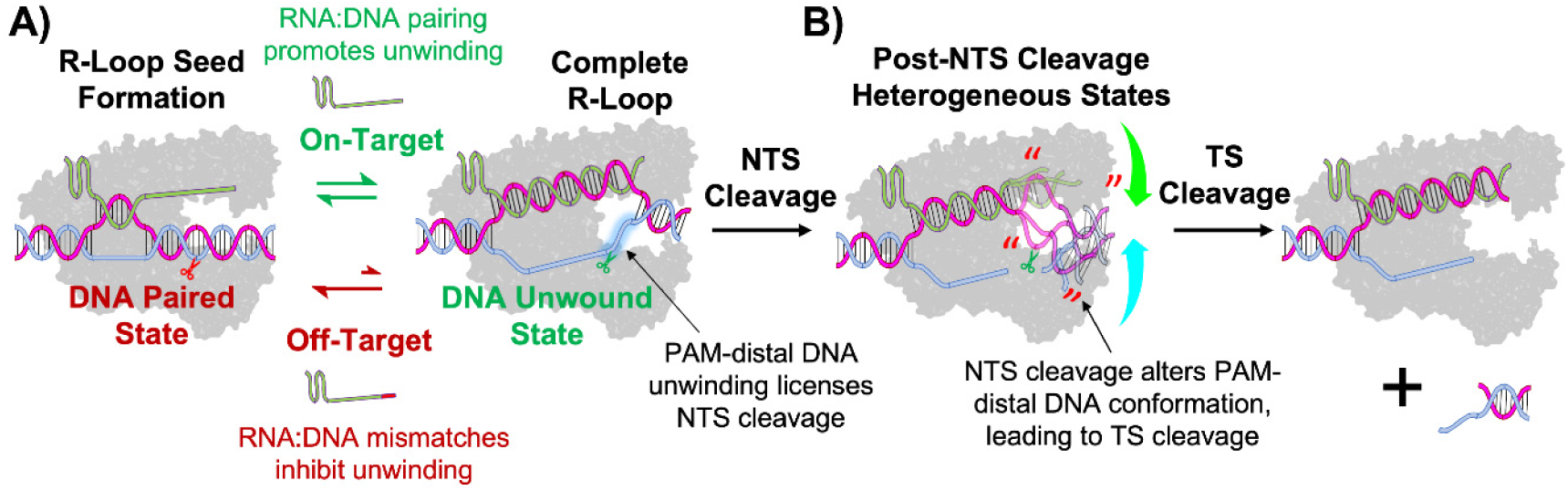
Proposed mechanism of Cas12a DNA target discrimination. (A) FnCas12a- induced unwinding of an intact DNA target adopts an inherent equilibrium between a duplex-like paired state and a DNA unwound state. This paired/unwound equilibrium serves as a checkpoint for the first-step of NTS cleavage, as on-target RNA/DNA pairing promotes PAM- distal DNA unwinding and licenses the target for the first-step NTS cleavage; while off-target mismatches trap the target at the duplex-like paired state and inhibit NTS cleavage. (B) First-step nicking of NTS alters DNA conformations at the TS cleavage site, allowing TS scission. The DNA paired/unwound equilibrium is no longer maintained to discriminate against RNA/DNA mismatches.

The presence of a duplex-like paired DNA state within the ternary complex in solution has significant functional implications. Cas12a possesses one nuclease domain belonging to the RuvC family, which acts as a single-stranded nuclease (10, 12, 13, 19). As such, occupancy of the unwound state, in which NTS is exposed as a single-strand, is a pre-requisite for NTS nicking, while occupancy at the duplex-like paired state would prevent NTS cleavage (nicking). Data presented here clearly show that PAM-distal RNA/DNA mismatches (Fig. 3) tunes the equilibrium between the duplex-like state and the unwound state, thereby regulating occupancy of the unwound state and controlling NTS nicking. The large shift of the equilibrium towards the paired state (Fig. 6A) with the PAM-distal RNA/DNA mismatch can be rationalized based on the weakening of the RNA/DNA hybrid interaction, which in turn favors the DNA TS/NTS pairing. As such, the paired-unwound equilibrium is utilized by Cas12a to interrogate the proper complementarity between the RNA guide and the DNA target. It serves as a checkpoint that licenses the first step of NTS nicking, and dictates the entire Cas12a reaction pathway.

Data presented further indicate that the paired/unwound mismatch sensing equilibrium is maintained only prior to the first step of NTS nicking (Fig. 3 and 4). A number of studies have revealed DNA conformational changes post NTS-nicking in Cas12a (24, 30, 31, 45). Consistent with these reports, data obtained here using pre-nicked DNA constructs show that DNA conformational changes occur prior to the second step of TS cleavage across both the R-loop flank (Fig. 4) and PAM-distal region of the TS (Fig. 5). The DEER measurements revealed that post-NTS cleavage, the PAM-Distal segment of the DNA samples a number of conformations (Fig. 4, 5, 6B) as opposed to occupying discrete unwound or duplex-like paired states (Fig. 3B and 4A). Furthermore, the data indicate that with the pre-nicked construct, the off-target complex likely maintains a certain ability to sample through the conformational states to achieve TS cleavage (Fig. 5A), thus drastically reducing discrimination (Fig. 5C and 5D). Overall, this leads to our proposed model that the paired/unwound equilibrium is a checkpoint placed prior to the first-step of the cleavage reaction pathway (Fig. 6).

The proposed paired/unwound equilibrium checkpoint in Cas12a is quite distinct from that of Cas9 and accounts for the observed differences in the mismatch sensitivity patterns (32, 33). Work has shown that Cas9 possess an “excess” unwinding ability prior to strand scission: It completely unwinds the protospacer with an on-target full-length RNA guide (41, 49); and supports a certain degree of unwinding either in the presence of PAM-distal RNA/DNA mismatches (38, 39) or with truncated guides that do not allow PAM-distal RNA/DNA pairing (41). Such “excess” unwinding allows the ternary complex to transition into the catalytically active conformation, leading to promiscuous DNA cleavage. This accounts for the observed high frequency occurrence of PAM-distal mismatches in the off-target pattern (32). On the other hand, with the FnCas12a studied here, the on-target complex does not excessively tilt the system towards unwinding, as a paired duplex-like state is observed to coexist with the fully unwound state (Fig 2B and D). Such an equilibrium enables the native Cas12a to place a more stringent requirement for matching between the RNA guide and the DNA, as each and all of the RNA/DNA pairings are required to maintain occupancy of the unwound state. Indeed, this conclusion is supported by a complete collapse of the unwound state in the off-target complex, which prevents NTS cleavage (Fig. 3). Coupled with the sequential cleavage model requiring nicking NTS first prior to TS cleavage, Cas12a processes an exclusive ability to distinguish PAM-distal mismatches, resulting in very low PAM-distal off-targets that sets it apart from Cas9 (33–35).

Expanding on prior work in Cas9 (41, 48, 49), data presented here showcase the unique ability of SDSL in deriving insight on the structure-dynamics-function relationship governing CRISPR target discrimination. The ability of SDSL and DEER to reveal ensemble population distributions in solution (59) enables the experimental observation of the Cas12a paired/unwound equilibrium that senses PAM-distal RNA/DNA mismatches (Fig. 2 and 3). The R5 label has a smaller size and a short linker, and therefore are much more intimately coupled to the DNA as compared to most fluorophores. This is critical for the experimental identification of a duplex-like state within the Cas12a bound DNAs (Fig. 2, 3, 4). While this study focuses on DNA conformations, work is on-going to apply SDSL to connect conformational changes in CRISPR protein domains to DNA unwinding and target discrimination.

In conclusion, this work harnesses the utility of SDSL to assess conformational equilibrium in solution and combines it with 2AP fluorescence spectroscopy and enzyme kinetics to characterize the mechanism of Cas12a target DNA unwinding and specificity. The data reveals an inherent DNA unwinding equilibrium that serves as a check-point for sensing PAM-distal mismatches prior to first-step cleavage of NTS. This sets a foundation for future exploration on enhancing specificity in Cas12a-based applications via manipulation of this unwinding equilibrium.

## DATA AVAILABILITY

All relevant data are included in the main text or the supplemental information.

## SUPPLEMENTARY DATA

Supplementary Data are available at NAR online.

## AUTHOR CONTRIBUTIONS

J.S. and P.Z.Q. designed the studies and wrote the manuscript; J.S. and A.A. prepared protein and RNA reagents; J.S. and K.G.L. carried out the enzyme kinetics measurements; J.S. carried out spin-labeling measurements and modelling; J.S. and W.J. carried out 2AP measurements.

## ACKNOWLEDGEMENTS

We thank X. Zhang, I. Schuster, D. Wu, and Y. Liu for comments on the manuscript.

## Funding

National Institution of General Medical Sciences (P.Z.Q., R01GM124413, R35GM145341, RR028992; J.S., T32-GM118289) and the Anton B. Burg Foundation. J.S. is a recipient of the Alfred E. Mann Innovation in Chemistry Doctoral Fellowship.

## CONFLICT OF INTEREST

The authors declare no Conflict of Interest.

## TOC

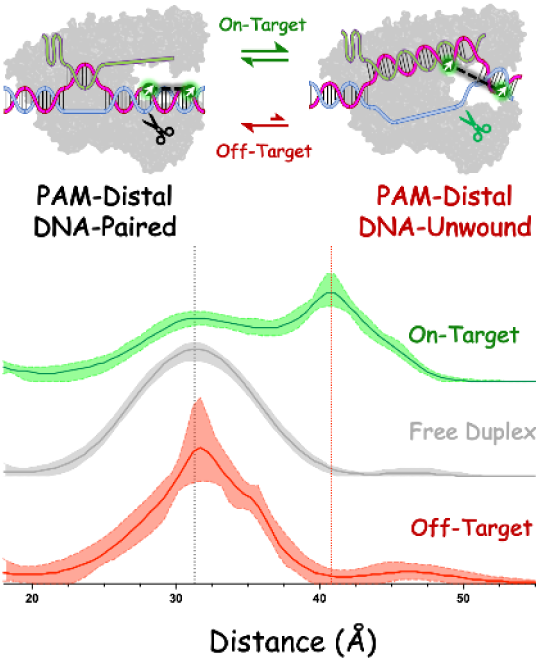

## Notes

### Competing Interest Statement

The authors have declared no competing interest.

