## Supplemental Information for "A DNA Unwinding Equilibrium Serves as a Checkpoint for CRISPR-Cas12a Target Discrimination"

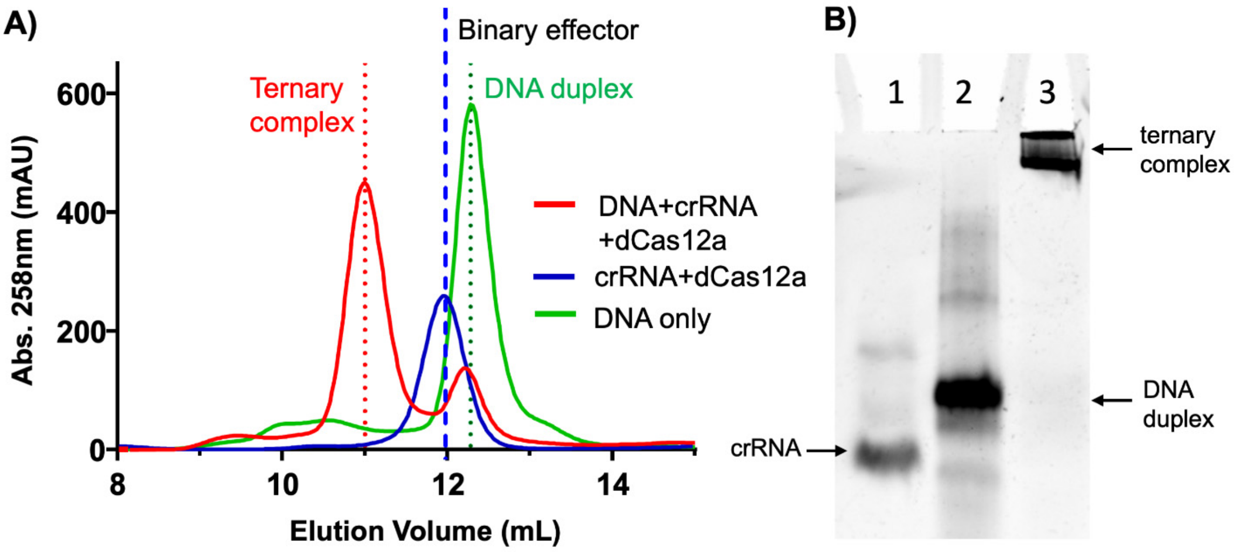

**Figure S1: Characterizations of dCas12a ternary complexes for spectroscopic measurements.** (A) A representative trace from size-exclusion-chromatography showing clear separation of the dCas12a bound DNA from the free duplex. (B) A native gel shift analysis of stability of the dCas12a ternary complex used in DEER measurements. Lanes 1, 2, and 3 are, respectively, crude P3 crRNA, free P3 DNA duplex, and a purified P3 on-target complex post DEER measurement. The samples were visualized via ethidium bromide staining. The DEER sample (lane 3) shows no observable free DNA duplex, indicating little dissociation of the DNA from the ternary complex under the conditions used to obtain DEER data.

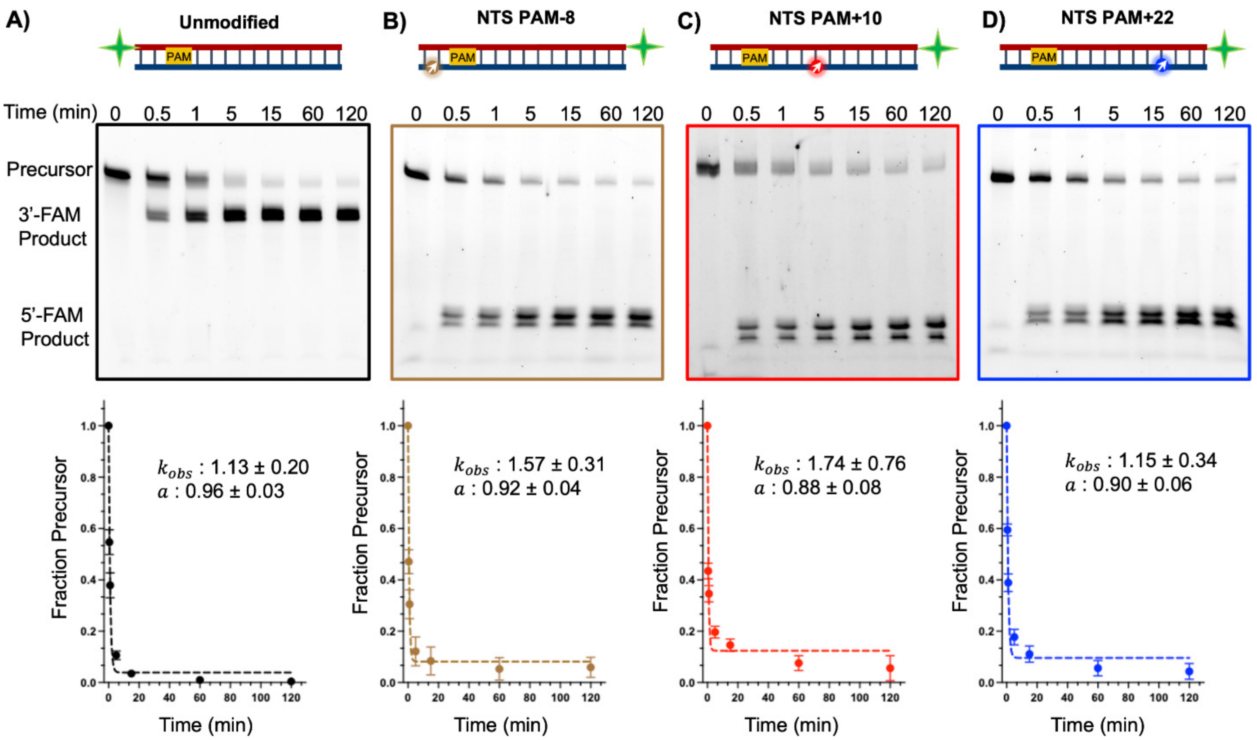

**Figure S2: Assessment of functional perturbations due to R5 labeling.** Cleavage time-course experiments (see Methods, main text) were performed with the unmodified P3 substrate (panel A) and P3 with an R5 label attached at PAM-8 (panel B), PAM+10 (panel C), and PAM+22 (panel D) of the NTS. In these measurements, the target-strand (TS) was monitored with a FAM label (indicated by the green star within the schematic), and the data were fit to eq.4 (main text) to obtain the TS cleavage rate constant ($k_{obs}$) (bottom panel), which reported on nicking of the non-target-strand followed by cleavage of the target-strand given the sequential cleavage mechanism of Cas12a. The measured $k_{obs}$ values of R5-labeled substrates are very similar to that of the unmodified substrate, indicating minimal perturbation due to R5 labeling at these positions. Note that in these measurement the multiple product bands indicate variability of Cas12a TS cleavage sites, which have been reported (1, 2). Furthermore, R5-labeled TS was not assessed by enzyme kinetic measurements due to the difficulty of generating the proper FAM- and R5-labeled TS strands. However, for DEER measurements with one or both R5 labels attached to TS, the resulting distance distribution profiles show a population matching what is predicted from the reported crystal structure (main text, Fig. 2B and D, Fig. 4A), thus supporting the notion that perturbations from R5 labels at these positions do not impact conclusions drawn.

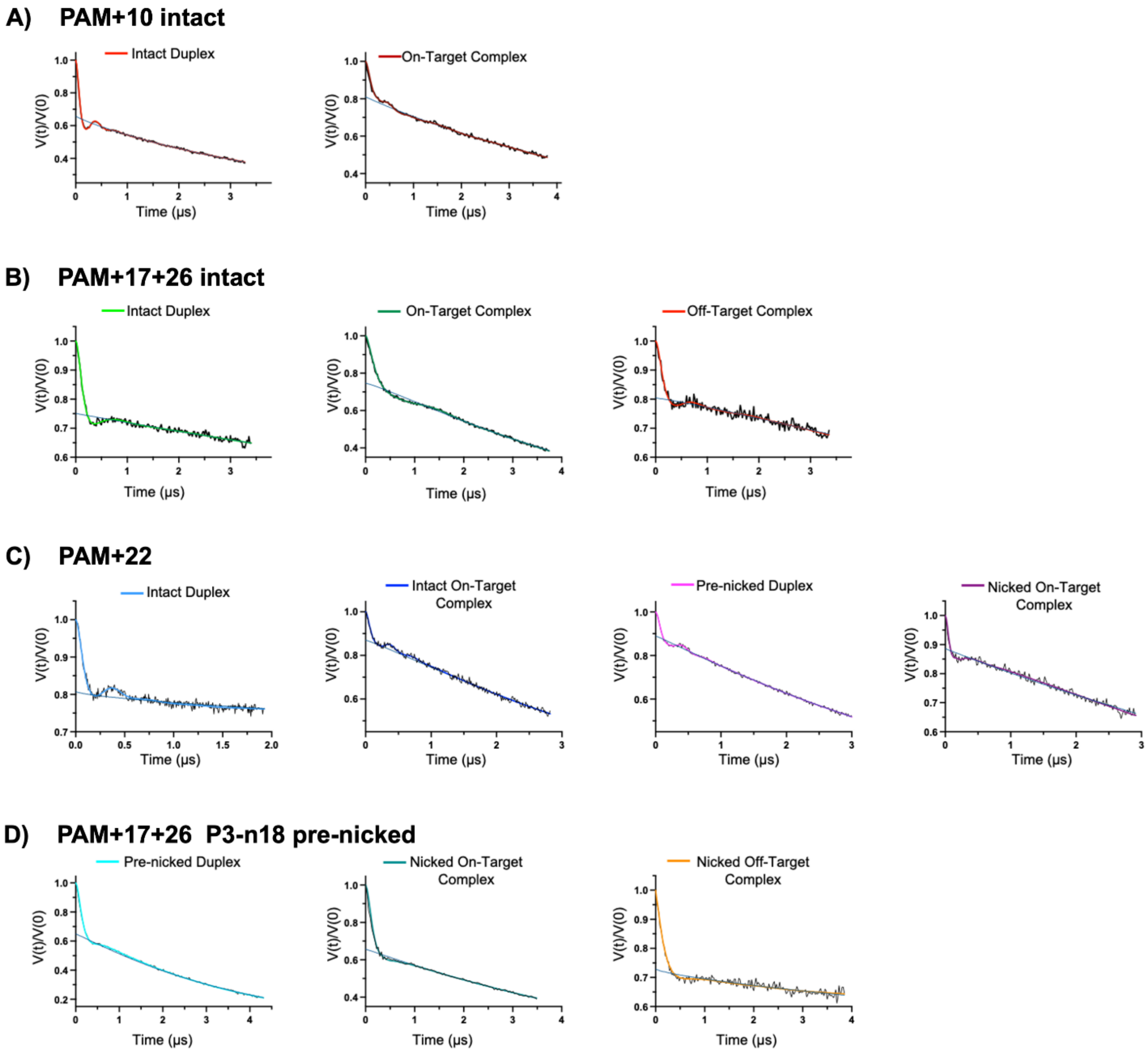

**Figure S3: Dipolar evolution traces from DEER measurements.** In each dataset, the raw dipolar evolution trace is shown in black, the background correction generated by the Consensus DEER program (3, 4) is shown in ocean blue, and the Consensus DEER fit (3, 4) is shown in color. (A) PAM+10 intact; (B) PAM+17+26 intact; (C) PAM+22; and (D) PAM+17+26 P3-n18.

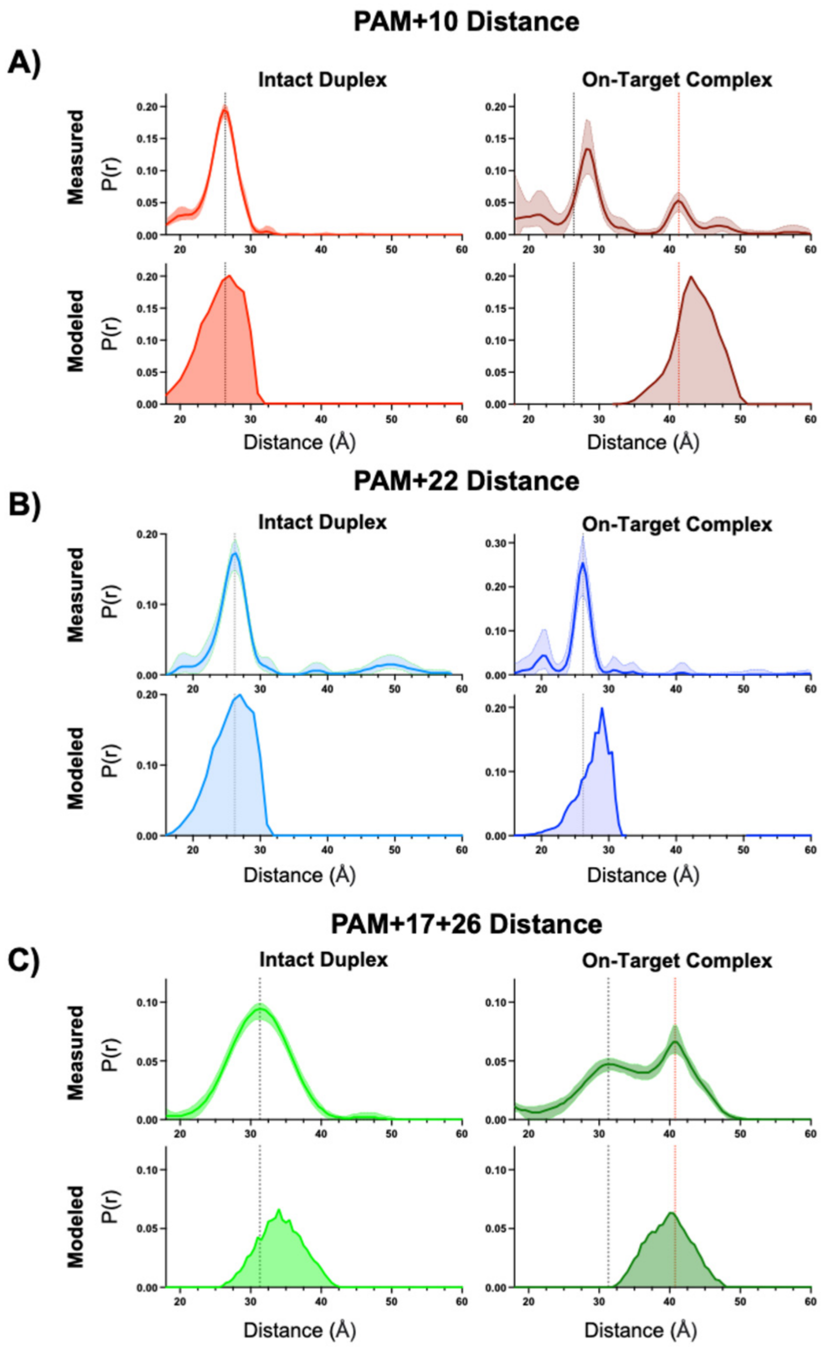

**Figure S4: Comparison of measured distance distributions with ALLNOX modeled distributions.** ALLNOX modeling was carried out as described in Methods in the main text. In each data set, the DEER measured distance distribution is shown on top, and the ALLNOX modeled distribution is shown at the bottom. To aid comparison, a black dashed line marks the measured maxima of the respective free duplex, and a red dashed line marks the measured maxima of the long-distance population in the respective complex samples. Overall, there is good agreement between the measured distributions of the free duplex with those modeled using a standard B-from DNA (see "Intact Duplex"), as well as between the measured long-distance populations with those modeled on a reported crystal structure (pdb id 6i1k) representing the DNA unwound state (see "On-Target Complex"). (A) PAM+10 construct. (B) PAM+22 construct. (C) PAM+17+26 construct.

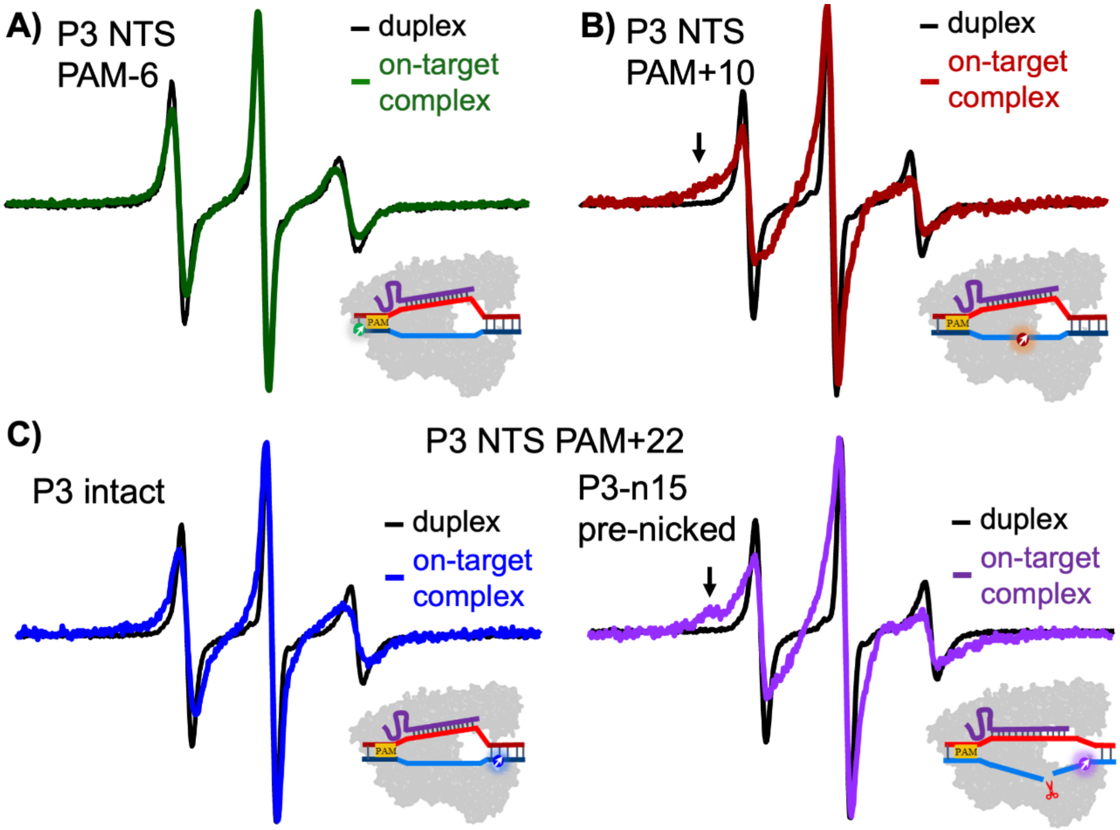

**Figure S5: Assessing the local environment at R5-labeling sites using cw-EPR spectroscopy.** Each dataset shows X-band cw-EPR spectra of a singly-R5-labeled duplex free in solution (black) and bound to an on-target dFnCas12a complex (colored). Spectra were acquired and processed as described in Methods in the main text. For comparison, the spectra were normalized to the same central-line amplitude. The lineshape of an X-band cw-EPR spectrum is dictated by the overall rotational dynamics of the R5 label, which is impacted by a combination of three factors (5) (i) the overall tumbling of the parent molecule; (ii) motion of the segment of the macromolecule at or near the labeling point; and (iii) torsional rotations about bonds that connect the nitroxide ring to the macromolecule; which are impact by steric contacts between the label and the parent molecule. As such, variations in the lineshape provide structural and dynamic information at the labeling site. (A) Spectra of an R5 label attached at the PAM-6 position of the NTS. Nearly identical spectra are observed between the free and bound DNA, indicating no change at the local DNA environment upon dCas12a binding. This is expected, as this position is beyond the PAM, and neither DNA unwinding nor contact(s) to the Cas12a effector is expected. (B) Spectra of an R5 attached at the PAM+10 position of NTS, which is located at the middle of the protospacer. The bound DNA spectrum shows immobilized components (indicated by the arrow) that are not presented in either the free DNA or the NTS-8 bound spectrum (Fig. S5A). These reflect new conformation(s) adopted by NTS upon unwinding, which restricts motions of a fraction of the labels due to additional contact(s) between the label and the dCas12a RNP. (C) Spectra of an R5 attached at the PAM+22 position of the NTS, which is directly across the TS cleavage site. With the intact duplex (left), the bound spectrum shows the same three-line characteristics as that of the free duplex, and no immobilized component is observed. This indicates that the label is free of contact with the effector, which is consistent with a reported crystal structure (pdb id 6i1k). With the P3-n15 pre-nicked duplex (right), the bound spectrum shows immobilized components (indicated by the arrow). This reflects alteration of label dynamics due to changes in DNA conformations upon first-step cleavage of the NTS.

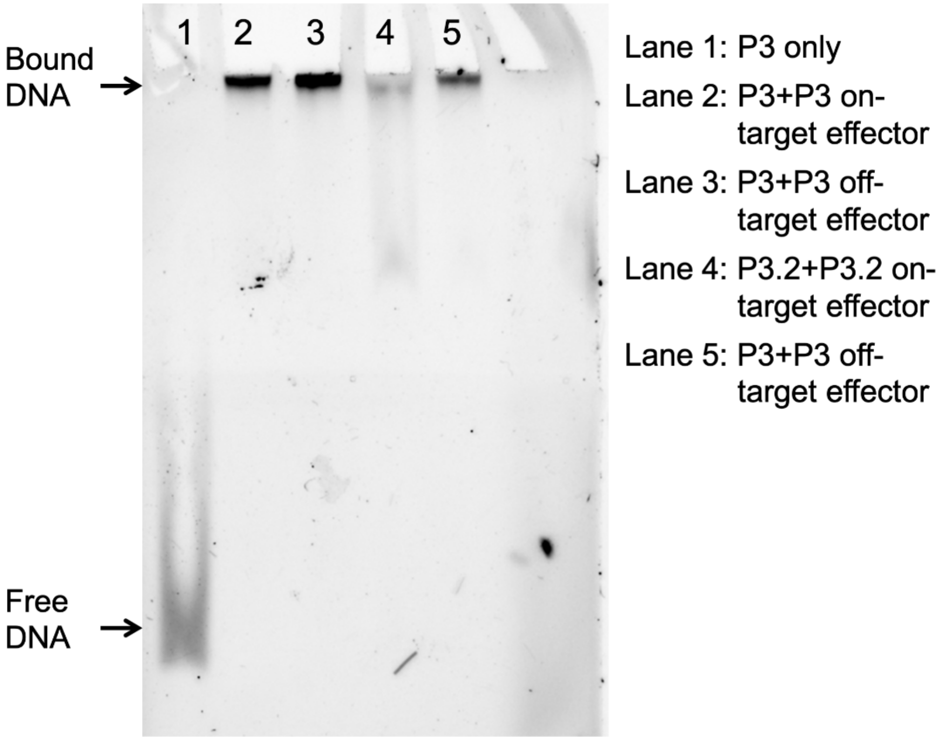

**Figure S6: Assessing substrate binding with on-target and off-target Cas12a effectors.** To assess substrate binding, FAM-labeled intact DNA duplex substrates were subjected to ternary complex assembly with a catalytically inactive effector formed by the dFnCas12a protein and either an on-target or an off-target crRNA. The concentrations of the reagents and the procedure for ternary assembly were the same as that reported in the cleavage time-course measurements (Methods, main text). The data show no observable free DNA duplex in the presence of either the P3 (lanes 2 and 3) or P3.2 effectors (lanes 4 and 5), indicating complete binding of substrates. This ensures that the cleavage kinetics data reported in this work was obtained under saturation condition. In addition, the lack of cleavage observed with the P3 off-target effector duplex (Fig. 3D, main text) was not due to substantially weakened binding.

**Table S1: Sequences of DNA constructs used in this study.**[a]

| | |
|---|---|
| **P3 Constructs**[b] | |
| **Target Strand** | |
| P3a | GGATGCCATGTGTGTGTGGAATGCCATGTGGGCTGTCTAAATTGAGCGAGTGGAA |
| P3a+10[c] | GGATGCCATGTGTGTGTGGAATGCCATG*TGGGCTGTCTAAATTGAGCGAGTGGAA |
| P3a+22[c] | GGATGCCATGTGTGTG*TGGAATGCCATGTGGGCTGTCTAAATTGAGCGAGTGGAA |
| P3a+17+26[c] | GGATGCCATGTG*TGTGTGGAA*TGCCATGTGGGCTGTCTAAATTGAGCGAGTGGAA |
| P3a-FAM-3'[d] | GGATGCCATGTGTGTGTGGAATGCCATGTGGGCTGTCTAAATTGAGCGAGTGGAA**/36-FAM/** |
| **Non-Target Strand** | |
| P3b[e] | TTCCACTCGCTCAATTTAGACAGCCCACATGGCATTCCACACACACATGGCATCC |
| P3b-5'FAM[d,e] | **/56-FAM/**TTCCACTCGCTCAATTTAGACAGCCCACATGGCATTCCACACACACATGGCATCC |
| P3b+10[c] | TTCCACTCGCTCAATTTAGACAGCCCA*CATGGCATTCCACACACACATGGCATCC |
| P3b+22[c] | TTCCACTCGCTCAATTTAGACAGCCCACATGGCATTCCA*CACACACATGGCATCC |
| P3b+23(2AP)[e,f] | TTCCACTCGCTCAATTTAGACAGCCCACATGGCATTCCAC**/i2AmPr/**CACACATGGCATCC |
| P3b-5’n18[g,i] | TTCCACTCGCTCAATTTAGACAGCCCACATGGCATT |
| P3b-3’n18[g,i] | CCACACACACATGCCATCC |
| P3b-5’n15[h,i] | TTCCACTCGCTCAATTTAGACAGCCCACATGG |
| P3b-3’n15[h,i] | CATTCCACACACACATGGCATCC |
| P3b-3’n15+22[c,h,i] | CATTCCA*CACACACATGGCATCC |
| P3b-3’n15+23(2AP)[e,h,i] | CATTCCAC**/i2AmPr/**CACACATGGCATCC |
| **P3.2 Constructs**[j] | |
| **Target Strand** | |
| P3a.2 | GGATGCCATGTGTGTGTGGTATGCCATGTGGGCTGTCTAAATTGAGCGAGTGGAA |
| **Non-Target Strand** | |
| P3b.2 | TTCCACTCGCTCAATTTAGACAGCCCACATGGCATACCACACACACATGGCATCC |
| P3b.2+18(2AP)[f] | TTCCACTCGCTCAATTTAGACAGCCCACATGGCAT**/i2AmPr/**CCACACACACATGGCATCC |

(a) Sequences shown as 5’->3’, with the PAM highlighted in yellow, and the protospacer underlined.
(b) Used with the corresponding P3 On-Target and P3 Off-Target crRNAs listed in Table S3.
(c) “*” indicates phosphorothioate modification(s).
(d) Fluorescein label at either the 5’ (“/56-FAM/”) or 3’ (“/36-FAM/”) terminus, used for cleavage assays.
(e) Hybridized with the appropriate P3a target-strand to form the intact P3 duplex.
(f) “/i2AmPr/” indicates a 2-amino-purine modification.
(g) Hybridized with the appropriate P3a target-strand to form the nicked P3-n18 duplex.
(h) Hybridized with the appropriate P3a target-strand to form the nicked P3-n15 duplex.
(i) FnCas12a has been shown to exhibit imprecise NTS cleavage and trimming activity between positions NTS+15 to NTS+18 (1). As such, while P3-n18 and P3-n15 place the nick at different NTS sites, they both mimic the post-NTS-cleavage DNA, and serve as the proper constructs to study progression towards TS cleavage by Cas12a.
(j) Used with the corresponding P3.2 On-Target and P3.2 Off-Target crRNAs listed in Table S3.

### Table S2: DNA templates used for T7 *in vitro* transcription to generate crRNA.[(a)]

| Template[(a)] | Sequence[(b)] |
|---|---|
| P3 On-Target | AAGTGGAATGCCATGTGGGCTGTCATCTACAACAGTAGAAATTCctatagtgagtcgtattag |
| P3 Off-Target[(c)] | AAGTCCTTTGCCATGTGGGCTGTCATCTACAACAGTAGAAATTCctatagtgagtcgtattag |
| P3.2 On-Target[(d)] | AAGTGGTATGCCATGTGGGCTGTCATCTACAACAGTAGAAATTCctatagtgagtcgtattag |
| P3.2 Off-Target[(d,e)] | AAGTCCATTGCCATGTGGGCTGTCATCTACAACAGTAGAAATTCctatagtgagtcgtattag |

(a) Templated named following the corresponding RNA nomenclature used in Table S3.
(b) Sequences shown as 5'->3'. Lower-case letters indicate the segment that hybridizes with a "T7 top strand" oligonucleotide (5'-ctaatacgactcactatag-3') to form the T7 promoter. Underlines indicate the segment to generate the RNA guide.
(c) Shown in red are alterations that creates four mismatches at PAM+17 to 20 with respect to the P3 DNA constructs.
(d) Yellow highlights sequence changes from the corresponding P3 templates.
(e) Shown in red are alterations that creates four mismatches at PAM+17 to 20 with respect to the P3.2 DNA constructs.

### Table S3: Sequences of crRNAs

| crRNA | Sequence[(a)] |
|---|---|
| P3 On-Target[(b)] | GGAAUUUCUACUGUUGUAGAUGACAGCCCACAUGGCAUUCCACUU |
| P3 Off-Target[(c)] | GGAAUUUCUACUGUUGUAGAUGACAGCCCACAUGGCA**AAGG**ACUU |
| P3.2 On-Target[(d)] | GGAAUUUCUACUGUUGUAGAUGACAGCCCACAUGGCAUACCACUU |
| P3.2 Off-Target[(e)] | GGAAUUUCUACUGUUGUAGAUGACAGCCCACAUGGCA**AUGG**ACUU |

(a) Sequences shown as 5'->3'. Underlines indicate the 20-nucleotide RNA guide that hybridizes with the target-strand of the protospacer. An extra four nucleotides are included at the 3' segment of the guide. It is known that these nucleotides do not interact with the protospacer [ref.]. They are included to prevent potential issues due 3' terminus heterogeneity in T7 generated RNA.
(b) Guide presents complete complementarity to PAM+1 to +20 positions of the P3 DNA constructs.
(c) Guide includes four mismatches (indicated in red) at PAM+17 to +20 of the P3 DNA constructs.
(d) Yellow highlights the one nucleotide change from the P3 On-Target RNA. Guide presents complete complementarity to PAM+1 to +20 positions of the P3.2 DNA constructs.
(e) Yellow highlights the one nucleotide change from the P3 Off-Target RNA. Guide includes four mismatches (indicated in red) at PAM+17 to +20 of the P3.2 DNA constructs.

**Table S4. Summary of DEER data quality and regularization parameters.**[(a)]

| | PAM+10 Intact | | PAM+22 Intact | | PAM+22 pre-Nicked | |
|---|---|---|---|---|---|---|
| | **Duplex** | **On target Complex** | **Duplex** | **On target Complex** | **Duplex** | **On target Complex** |
| Modulation Depth (%) | 16.1% | 18.4 | 19% | 10.1% | 10.9 | 11.70% |
| SNR (WRT) | 18 | 29.1 | 26.8 | 23 | 22.3 | 13.8 |
| alpha-value-DEERNET (CDA) | 1 | 0.45 | 0.7 | 0.12 | 0.14 | 0.56 |
| alpha-value Tikhonov | 3.98 | 0.32 | 7.94 | 2 | 1.58 | 1.58 |
| CDA overlap | 82.8% | 81.7 | 88.7 | 92.6% | 91.8% | 86.7 |

| | PAM+17+26 Intact | | | PAM+17+26 pre-Nicked | | |
|---|---|---|---|---|---|---|
| | **Duplex** | **On Target Complex** | **Off Target Complex** | **Duplex** | **On Target Complex** | **Off Target Complex** |
| Modulation Depth (%) | 24.7 | 25.4 | 19.2 | 34.8 | 37 | 26.3 |
| SNR (WRT) | 31.9 | 49.4 | 16 | 82.8 | 77.9 | 26.5 |
| alpha-value-DEERNET (CDA) | 3.54 | 1.25 | 1.25 | 2.81 | 7.07 | 7.92 |
| alpha-value Tikhonov | 10 | 19.95 | 5.01 | 7.94 | 10 | 15.85 |
| CDA overlap | 96.2 | 96 | 89.6 | 92.5 | 93.5 | 93 |

(a) Details of these parameters can be found in reference (6), which describes standards established by the community for assessing DEER data and the associated distance distributions. Based on these standards presented in reference (6), DEER data reported in this work are sufficient to support the analysis presented and conclusions drawn.

**Table S5. Summary of 2-amino-purine data.**

| 2AP Position | Construct | $F_{368}/A_{260}$ (x $10^5$ a.u.)[(a)] | ε ($M^{-1}$ $cm^{-1}$) | ratio(φ)[(b)] |
|---|---|---|---|---|
| NTS+18[(c)] | P3.2 unbound duplex | 1.16 ± 0.057 | 869921 | 1 ± 0.10 |
| | P3.2 on-target complex | 2.19 ± 0.249 | 1319921 | 2.87 ± 0.47 |
| | P3.2 off-target complex | 1.09 ± 0.150 | 1319921 | 1.42 ± 0.27 |
| NTS+23[(d)] | P3 unbound duplex | 7.28 ± 0.008 | 869921 | 1 ± 0.02 |
| | P3 on-target complex | 4.54 ± 0.016 | 1319921 | 0.95 ± 0.04 |
| | P3-15nick unbound duplex | 2.33 ± 0.564 | 869921 | 1 ± 0.34 |
| | P3-15nick on-target complex | 16.96 ± 0.159 | 1319921 | 4.80 ± 0.88 |

(a) Average and standard deviation (std) obtained from at least three repeats. Reported as "average ± std" in arbitrary unit (a.u.).
(b) Computed according to eq. (2) in the main text. Reported as "average ± std" with std obtained from propagation of "errors", i.e. std obtained for the corresponding $F_{368}/A_{260}$ measurements.
(c) Data presented in Figure 3C of main text.
(d) Data presented in Figure 4C of main text.
